## Supplemental Materials for "What Happens After an Error?"

Supplementary Information

| **% of Perseverative Errors (out of all errors)** | | | | | |
| --- | --- | --- | --- | --- | --- |
| *Processing Time (in ms)* | *Previous trial* | *Experiment 1* | *Experiment 2* | *Experiment 3* | *Experiment 4* |
| *0-500* | *Incorrect* | 33.96 | *31.05* | *39.38* | *22.02* |
| *501-1000* | *Incorrect* | *37.68* | *47.93* | *40.77* | *41.67* |
| *1001-1500* | *Incorrect* | *36.07* | *72.41* | *41.41* | *45.21* |
| *1501-2000* | *Incorrect* | *40.38* | *72.73* | *51.95* | *32.31* |
| *0-500* | *Correct* | *37.41* | *36.67* | *37.30* | *26.00* |
| *501-1000* | *Correct* | *28.26* | *30.16* | *20.36* | *17.34* |
| *1001-1500* | *Correct* | *15.73* | *30.00* | *20.15* | *10.42* |
| *1501-2000* | *Correct* | *22.45* | *25.00* | *14.29* | *9.52* |

Supplementary Table 1. Proportion of errors that are due to perseveration at different processing time (PT) bins. Across all experiments, participants have some initial tendency to perseverate after both correct and incorrect trials. However, errors that occur on post-error trials are much more likely to be perseverative errors, especially at the longer PTs.

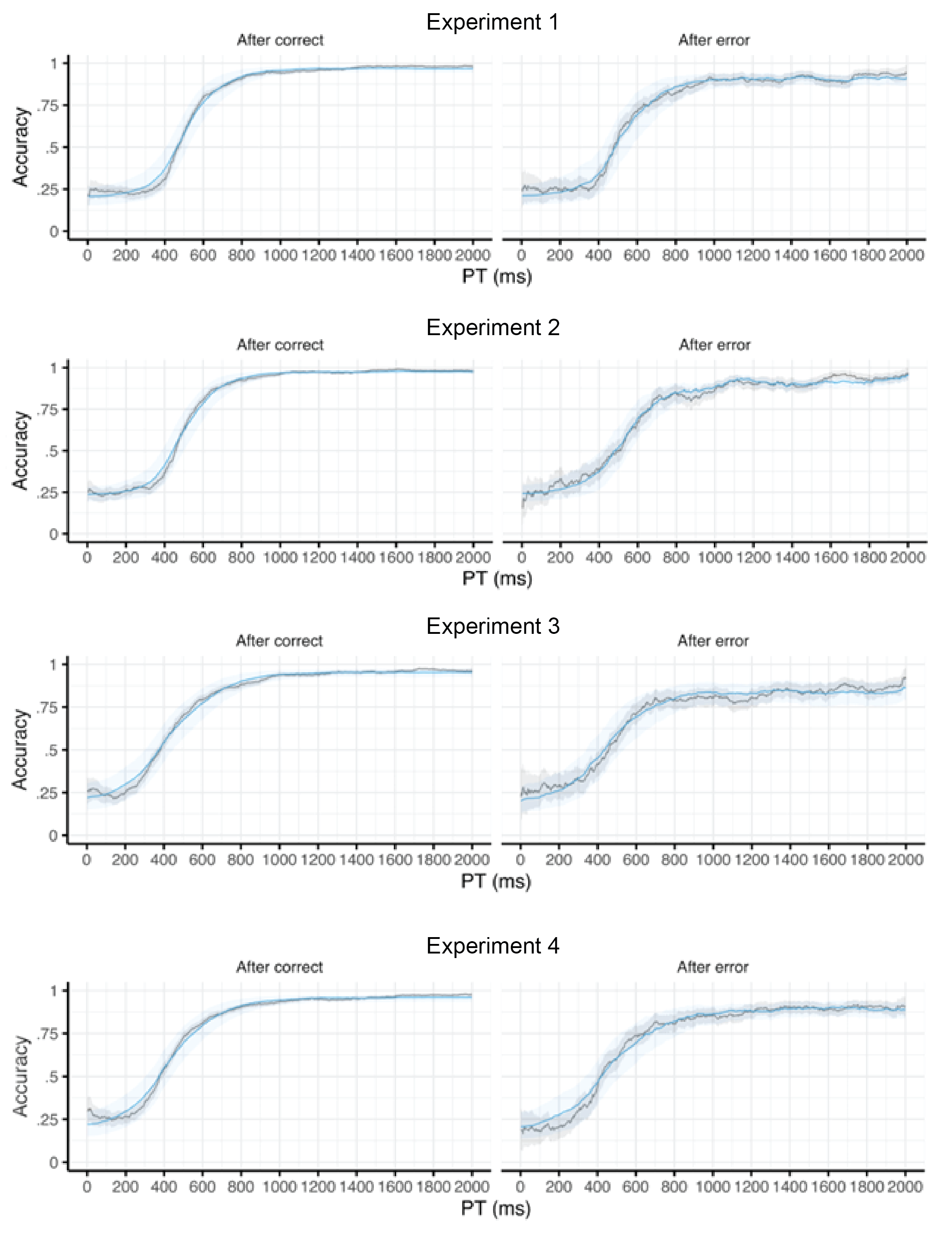
Supplementary Figure 1. Posterior predicted accuracy of fitted models in Experiments 1-4. Gray lines depict smoothed accuracy rate across all participants as a function of PT. Blue lines depict model-predicted accuracy rate using the median parameter estimates after model fitting. Note the close fit between the model’s predictions and participant behavior, especially at the later PTs that are primarily governed by the β parameter in our model that shows the primary post-error effects.

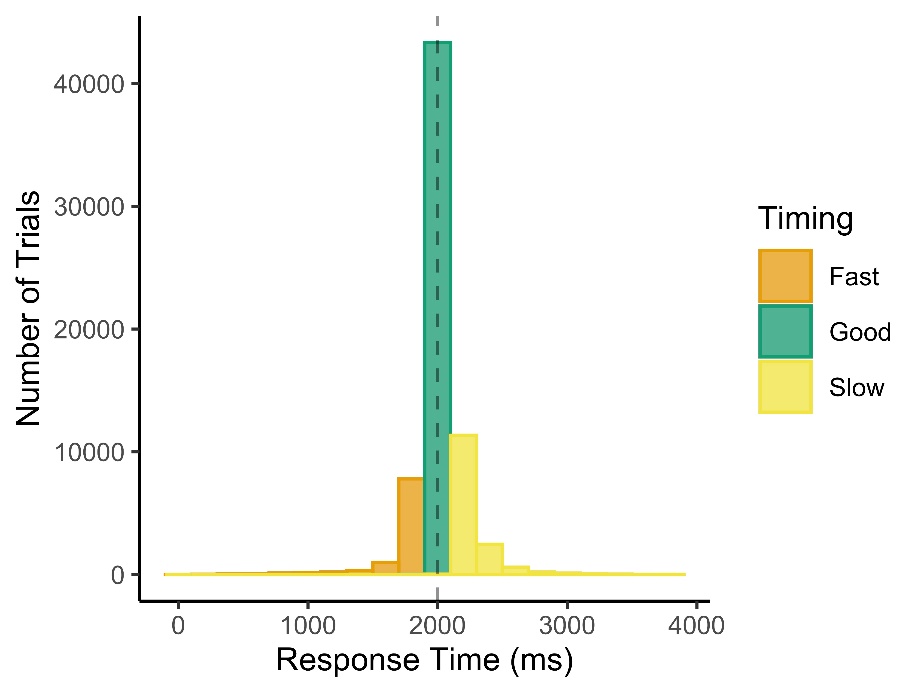

Supplementary Figure 2. Histogram of response times across all experiments. Participants were asked to make a response at 2000 ms (+/- 100 ms) following the onset of each trial. The vast majority of responses were on-time and “Good”. Although timing errors were approximately normally distributed, there was a slight skew toward “too slow” errors where the participants’ responses came after 2100 ms from the start of the trial.

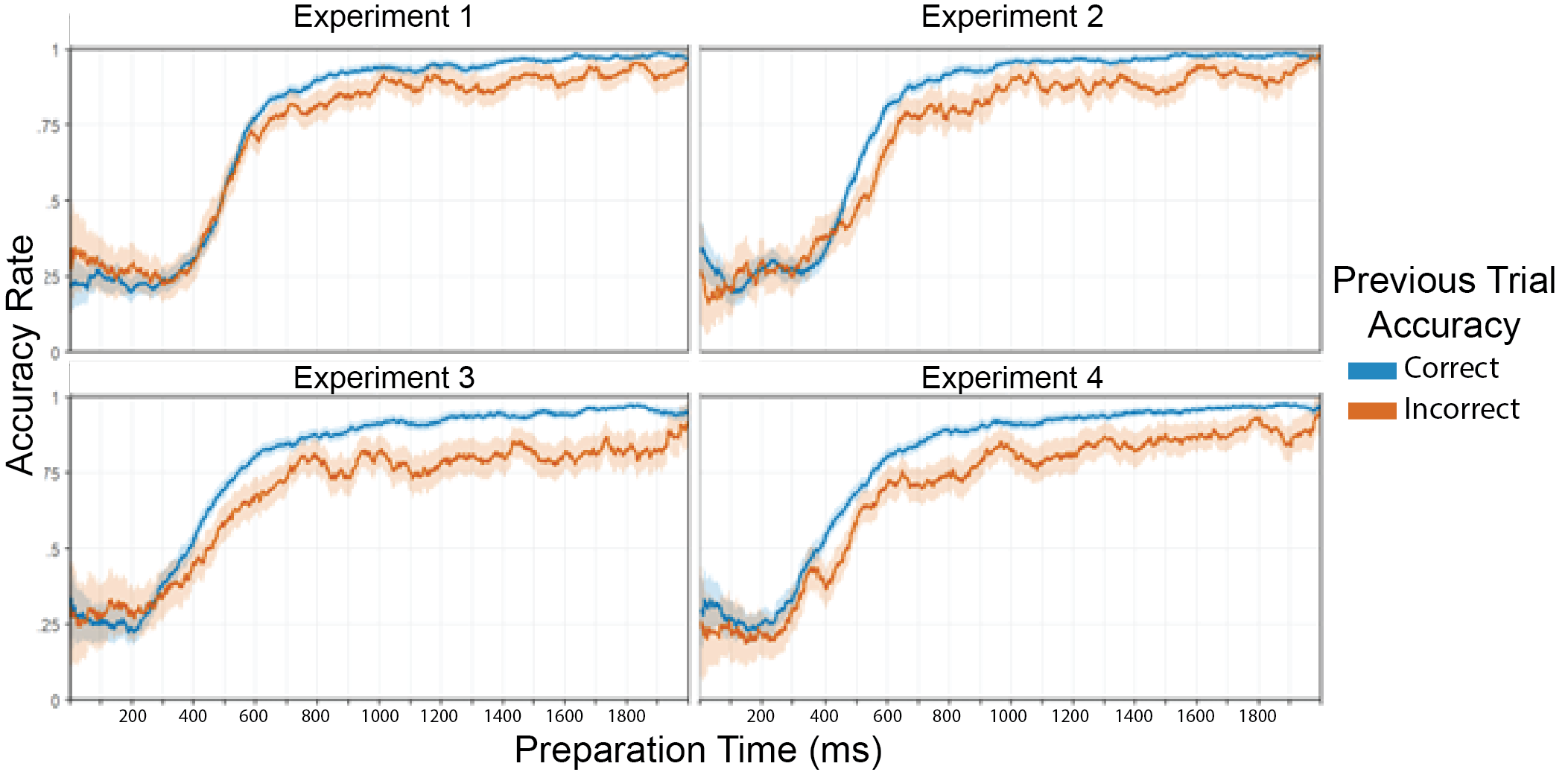

Supplementary Figure 3. Behavioral results with no timing (i.e. too fast/too slow) exclusions. Only trials on which the participants’ responded “on-time” within 100 ms of the go signal were included in the analyses presented in the main body of the manuscript. The data in the above figure includes all trials for each experiment regardless of the timing of the responses and shows a similar pattern of results.

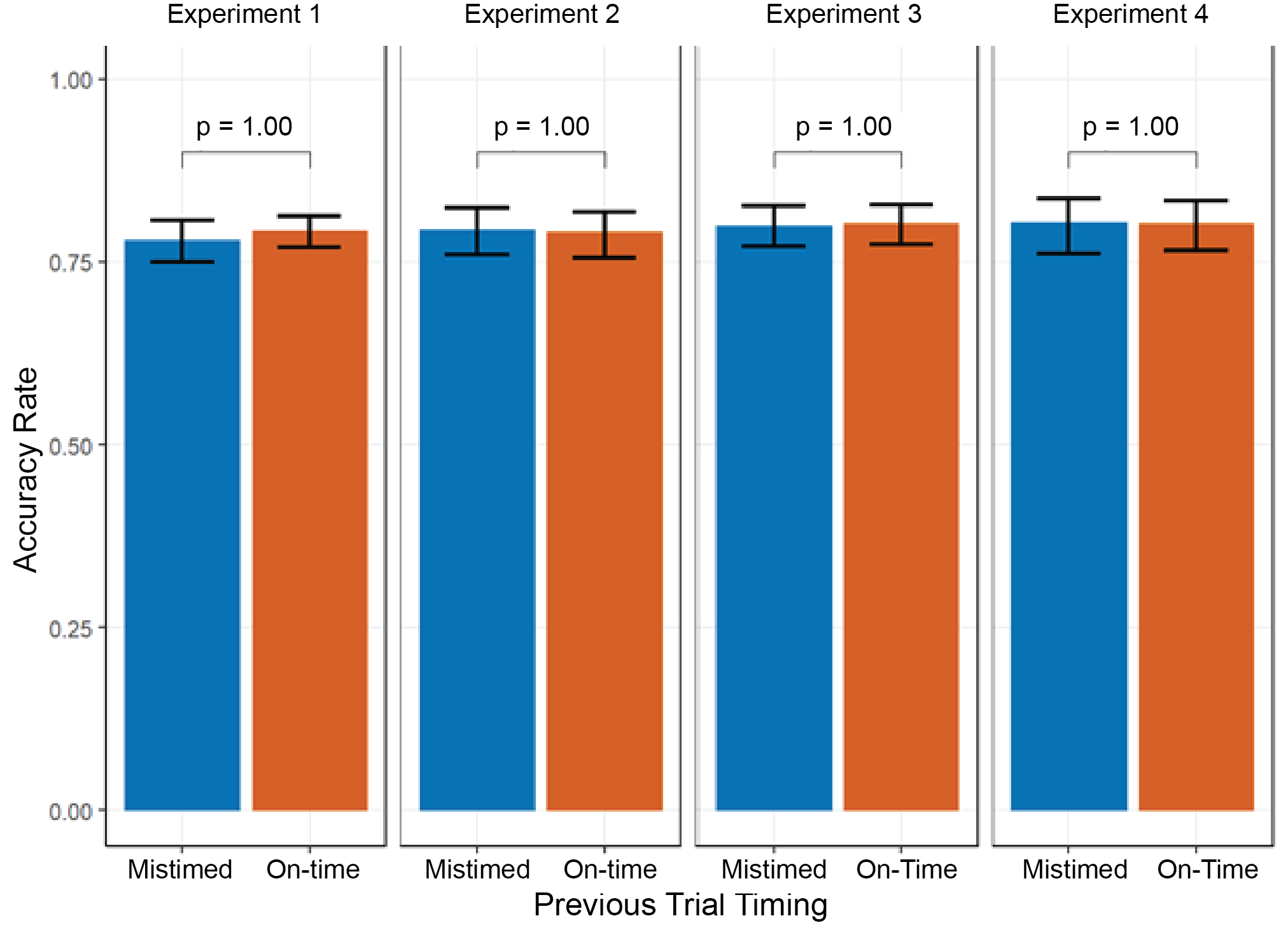

Supplementary Figure 4. Response accuracy following on-time and mis-timed trials. Across all four experiments, we did not find evidence that timing errors affected response accuracy on the subsequent trial. (P-values reflect Bonferroni-corrected significance values from paired t-tests for each experiment.).)

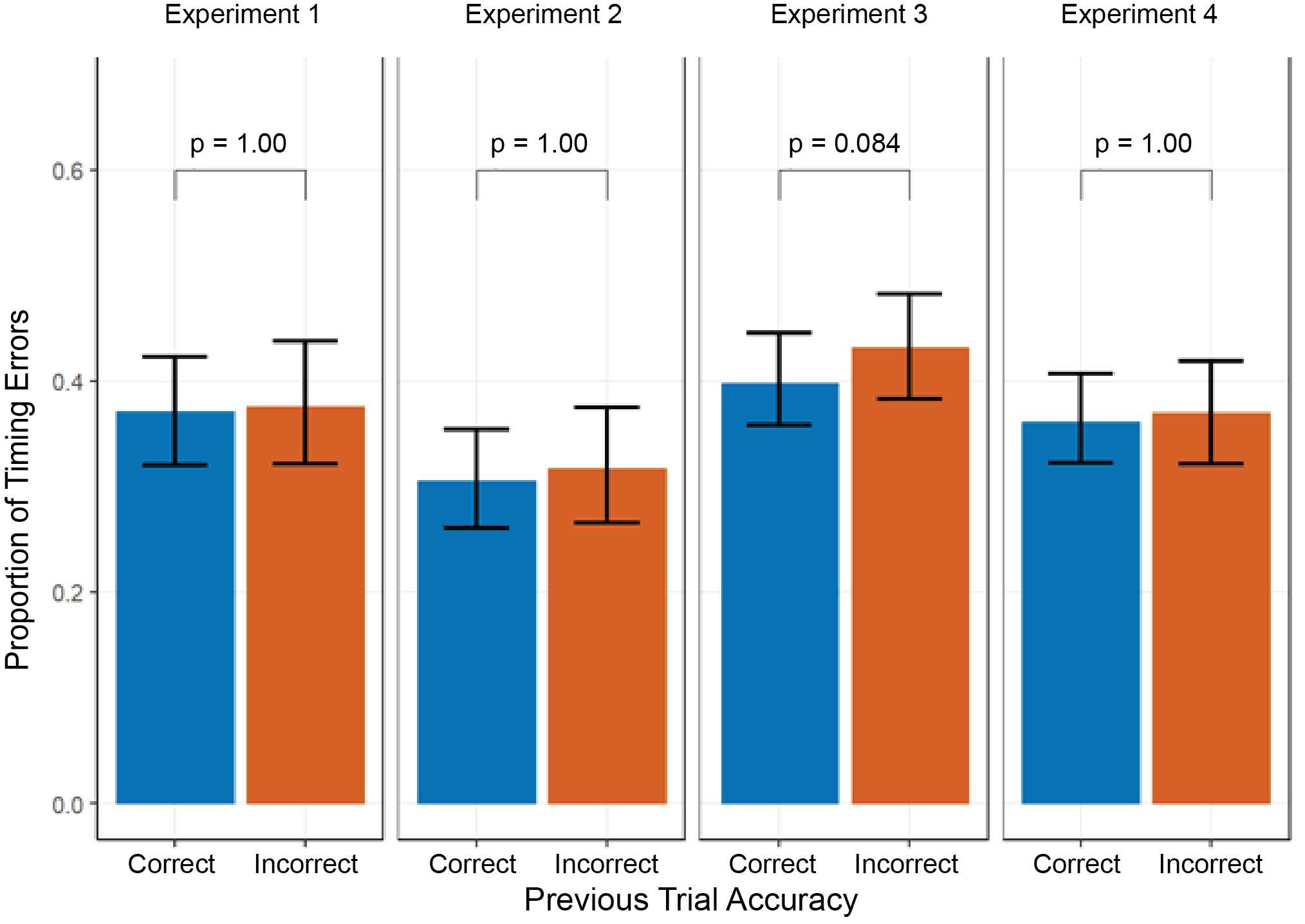

Supplementary Figure 5. Proportion of timing errors following correct and incorrect responses. Participants do not appear to be substantially more likely to mistime their responses on trials following an error. (P-values reflect Bonferroni-corrected significance values from paired t-tests for each experiment.)
